# Demography-environment relationships improve mechanistic understanding of range dynamics under climate change

**DOI:** 10.1101/2022.09.23.509134

**Authors:** Anne-Kathleen Malchow, Florian Hartig, Jette Reeg, Marc Kéry, Damaris Zurell

## Abstract

Species responses to climate change are widely detected as range and abundance changes. To better explain and predict them, we need a mechanistic understanding of how the underlying demographic processes are shaped by climatic conditions. We built spatially-explicit, process-based models for eight Swiss breeding bird populations. They jointly consider dispersal, population dynamics and the climate-dependence of three demographic processes - juvenile survival, adult survival and fecundity. The models were calibrated to two-decade abundance time-series in a Bayesian framework. We assessed goodness-of-fit and discriminatory power of the models with different metrics, indicating fair to excellent model fit. The most influential climatic predictors for population performance were the mean breeding-season temperature and the total winter precipitation. Maps of overall growth rate highlighted demographically suitable areas. Further, benefits from contemporary climate change were detected for typical mountain birds, whereas lowland birds were adversely affected. Embedding generic process-based models in a solid statistical framework improves our mechanistic understanding of range dynamics and allows disentangling the underlying abiotic and biotic processes. For future research, we advocate a stronger integration of experimental and empirical measurements and more detailed predictors in order to generate precise insights into the processes by which climate affects populations.

## 1 Introduction

Changing climatic conditions are impacting natural systems around the globe, causing rapid biodiversity changes [1, 2, 3]. Two of the most striking and commonly discussed impacts are distribution shifts [4] and changes in population abundances [5]. These range dynamics result from an interplay of key ecological processes such as local population dynamics and dispersal, which are widely considered to be influenced by the environment [6, 7]. An improved, model-based understanding of how climate affects range and population dynamics through these key processes is needed to better explain and predict the observed responses [8, 9]. Such insights can further form a quantitative, science-guided basis for deriving effective conservation measures to mitigate biodiversity loss [10].

A process-based approach to species distribution modelling has been suggested repeatedly, going beyond purely correlative models [11, 12, 13, 14]. It is expected that process-based models can provide more reliable predictions under changing conditions by explicitly including causal eco-evolutionary mechanisms [15, 6] and allowing for the representation of transient dynamics. Progress towards this goal was made with hybrid models that couple a phenomenological habitat model with processes like population dynamics and dispersal [16, 17]. However, hybrid models do not allow to establish an explicit link between demographic processes and the environment. Instead, they assume a relationship with a composite quantity, the suitability, which is often derived from correlative models and combines multiple land cover and climate variables. Yet, the relation of such suitability measures to both abundance [18] and to growth rates [19] has been questioned. This limitation can be overcome by considering explicit responses of processes like demography to environmental predictors. (See, e.g. Schurr et al. [20], who propose a spatially-explicit process-based model that considers parametric demography-environment relationships together with mechanistic dispersal effects.)

Direct measurements of demography-environment relationships can be obtained by measuring demographic rates over an environmental gradient, which requires large-scale and well-designed monitoring schemes [21]. This has been done for a number of plants [22, 23], but only rarely for animal species, for example fish (brown trout, [24]), or birds (North-American forest birds, [25], and Arctic sea ducks, [26]). As an alternative to direct measurements, Pagel & Schurr [27] suggested an inverse modelling approach that simultaneously estimates the demography-environment relationships and all other process parameters from empirical data. In the original formulation, Pagel & Schurr [27] assumed a logistic growth (as the Ricker model) to describe local population dynamics. To allow for more complex life histories, however, this approach needs to be extended to a population model that considers different demographic sub-processes, such as survival and fecundity, as well as their respective environmental responses [28].

The complexity required to model such detailed population dynamics can be accommodated in an individual-based model (IBM, [29, 30]) that implements algorithmic representations of all processes considered at the scale of individuals. Here, we built single-species IBMs for nine Swiss breeding bird populations and explicitly model the demographic sub-processes of juvenile survival, adult survival and fecundity together with dispersal. We calibrate the models in a Bayesian framework to simultaneously infer the parameters of the demography-environment relationships, density-dependence and dispersal of each species from survey data covering two decades of climate change. Mountainous regions like the Swiss Alps are particularly susceptible to current climate change, and altered temperature and precipitation patterns are already being observed [31]. In Switzerland, the detected trend in air temperature increase over five decades (1959-2008) reached 0.35 K/decade, which equates to about 1.6 times the northern hemispheric warming rate [32]. We therefore expect that bird species that live in such mountainous regions have already been responding to climate change during the past decades.

The models are built using the individual-based modelling platform RangeShiftR [33]. Each IBM is based on species-specific habitat maps derived from main habitat preferences, which were obtained from published species trait tables. Different climate layers that summarise key climatic variables during decisive periods of the year directly inform the spatio-temporal variation in demographic rates via parametric relationships. Since we considered only climatic predictors, we henceforth refer to these relationships as demography-climate relationships (DCRs). We examine the fitted DCRs for patterns across species and point out potential limitations in their interpretation, which originate from our data-driven calibration approach. To evaluate the DCRs, we map the demographic rates (juvenile survival, adult survival, fecundity) as well as the resulting local growth rates across Switzerland. Based on the calibrated model, we assess the impact of two decades of contemporary climate change on both the growth rate as well as on total abundance. Such insights can facilitate the communication of severe consequences of climate change as well as the design of potential mitigation measures, targeting the specific demographic processes that are most impacted. Our approach is applicable to any population for which spatio-temporal abundance data are available. It can be flexibly extended to allow more detailed conclusions by incorporating more complex DCRs and using more fine-grained predictors.

## 2 Materials and Methods

### 2.1 Study area & data

Switzerland is located in the Alps and thus features strong elevational gradients. The warming rates due to climate change show high spatial and seasonal variance, with their peak in summer at 0.46 K/decade and large values in the lowlands during autumn and in middle and high elevations in spring [32]. To describe the climatic variation over this landscape, we used bioclimatic data from CHELSA v2.1 [34, 35]. It provides monthly means of daily mean, minimum and maximum temperatures, as well as total precipitation for the years 1901-2019 at a spatial resolution of 30 arcsec (≈ 1 km). These climate layers were averaged over several months for the breeding season (April-July), autumn (September-November) and winter (December-February), and standardised using the mean and standard deviation over the considered set of years (1997-2019). We further used land cover data from the CORINE project [36] for 2000, 2006, 2012, and 2018 at a resolution of 100 m to inform species-specific maps of suitable habitat (based on published information on habitat preferences, see below). Parameter inference was based on data from the standardised and designed Swiss breeding bird survey (MHB), which produces yearly time series from 1999 to 2019 at 267 representatively selected sites. Each site comprises a 1 km^2^ square in which the number of breeding pairs was counted during two to three repeat surveys per year using the so-called simplified territory mapping [37]. To develop models of initial abundance, we further used Atlas data of the period 1993-96 Schmid et al. [38] that provides a snapshot of counts at 2318 sites randomly distributed across Switzerland.

### 2.2 Model building and fitting

We selected a list of bird species according to a set of common characteristics, which allowed us to use the same model structure for all. We chose passerines, that use forests, shrubs and mountainous regions as their main habitats, are sedentary or short-distance migrants, have a similar life history in which 1-year-olds can be considered mature and able to reproduce, and show changes in spatial abundance between the two Atlas periods of 1993-96 and 2013-16. Focusing on forest and upland species meant that effects of land use change such as intensification of agriculture are less likely to contribute substantially to population dynamics [39]. By excluding long-distance migrants we could assume that local winter conditions are meaningful predictors of demographic rates. These criteria allowed us to isolate the effects of climate on the population dynamics as much as possible. Further constraints were imposed in order to keep the parameter calibration feasible: The relatively simple life history was chosen to limit the number of demographic parameters and a minimum number of 170 non-zero counts in the MHB data was required. Overall, the criteria related to the species’ ecology and technical aspects of the parameterisation were fulfilled for nine species: Eurasian bullfinch, european crested tit, eurasian treecreeper, eurasian nuthatch, dunnock, goldcrest, common linnet, ring ouzel, and alpine accentor.

Despite their common characteristics, we expected that the demography of the species will respond differently to climate variation [40, 41]. To understand these responses, we first derived simple habitat maps for each species. For this, we determined a set of habitat classes from the habitat preferences given in Storchová & Hořák [42], that coarsely delineate the typical habitat of each species. These habitat classes were mapped to CORINE land cover classes (see Table A.2), yielding binary habitat maps at the resolution of 100 m. Then, the maps were spatially aggregated to a resolution of 1 km by calculating the suitability index *h_i_*. It counts the number of (100-m)^2^-habitat cells located within each 1-km^2^-landscape cell *i,* thus reaching from 0 to 100. A 10-km buffer around the Swiss border was retained to reduce boundary effects.

The derived habitat maps at a resolution of 1 km were then used within the RangeShiftR IBM [33] to simulate the population and dispersal dynamics of each species on a regularly gridded landscape of Switzerland. We modelled population dynamics with a female-only, 2-stage model comprising the stages “juvenile” and “adult”, with a transition time of one year between the two stages. The local population dynamics of this model are characterised by three demographic rates: juvenile and adult survival probability, *s_j_* and *s_a_,* and fecundity *ρ*. In our model, survival probability describes the annual mortality that primarily occurs during winter and additionally during autumn for juveniles. The modelled fecundity *ρ* includes all contributions to juvenile survival that occur before the juveniles are independent. Therefore, also nestling mortality or early juvenile mortality are included in *ρ*. Fecundity was further assumed to be density-dependent, decreasing exponentially with the ratio *n_i_b_i_* of local population density *n_i_* and local resource availability 1/*b_i_*. Resource availability was expressed as the overall strength of demographic density-dependence 1/*b*, modulated by the local suitability *h_i_*, as *b_i_* = *b/h_i_* * 100. Natal dispersal was modelled with a constant emigration probability *p_e_* for each juvenile to start a dispersal event. To identify the destination cell, an exponential dispersal kernel with mean dispersal distance *d* and uniformly distributed direction was evaluated individually. If the destination was a suitable cell, the juvenile settled in it. Else, if one of the eight directly neighbouring cells was suitable, the individual settled there and otherwise died. Hence, juveniles suffered a dispersal mortality additional to the annual mortality that was denoted by *s_j_*. The order of processes in each simulation year was first reproduction, then dispersal, and lastly survival. We used stochastic initial conditions for each model run by drawing from a auto-regressive distribution model [43] that was fitted to the Atlas data of the period 1993-96.

To formulate the DCRs, each demographic rate was modelled using a generalised linear model, with a logarithmic link function for fecundity *ρ* and a logistic link for survival (*s_j_* and *s_a_*). The predictors included mean or minimum temperature and total precipitation, averaged over the period of the year that was considered most relevant for the respective process (Table 1): Fecundity depended on the conditions during the breeding period (April-Juli); survival probabilities depended primarily on the winter conditions (December-February), with juvenile survival additionally affected by conditions in autumn (September-November) when juveniles are independent and are known to suffer from high mortality. All predictors were included with linear and quadratic polynomial effects, apart from minimum winter temperature, which was only included in linear form. Minimum winter temperature was highly correlated with autumn temperature (≈ 0.76), and was therefore excluded as predictor for *s_j_* (but still included for *s_a_*). Therefore, the DCR of temperature on juvenile survival should be interpreted as the combined effects from autumn and winter temperature. To summarise, the climatic variables determined all three demographic rates while the habitat suitability only influenced the density-dependence effect on fecundity.

**Table 1:** Aggregated climatic predictors and their responses as modelled by the demographyclimate relationships. The climatic variables of mean temperature, minimum temperature, and total precipitation were averaged over a relevant season and, up to the given order, related to a demographic rate as response variable. ()* - this effect was originally considered, but excluded due to high collinearity of *T_wn_* with *T_at_.*

| climatic predictor | season | abbr. | order | response |
| --- | --- | --- | --- | --- |
| mean temperature during breeding season | April-Juli | $T_{br}$ | 2 | $\rho$ |
| total precipitation during breeding season | April-Juli | $P_{br}$ | 2 | $\rho$ |
| mean temperature during autumn | Sept.-Nov. | $T_{at}$ | 2 | $s_j$ |
| minimum temperature during winter | Dec.-Feb. | $T_{wn}$ | 1 | $(s_j)^*, s_a$ |
| total precipitation during winter | Dec.-Feb. | $P_{wn}$ | 2 | $s_j, s_a$ |

We estimated the parameters of each DCR simultaneously with all other model parameters (1/*b, p_e_*, 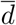) from MHB data using Bayesian inference [44]. As priors for the climate-independent parameters and the intercepts of the DCRs we assumed Gaussian distributions with means and standard deviations derived from the demographic traits provided by Storchová & Hořák [42] (Table A.1). For the DCR coefficients, we used mildly regularising truncated normal priors with *β_m,n_* = 0 ± 1 and truncations at [-5, 5]. For the counts, we followed the usual approach and assumed a negative-binomial likelihood. Its dispersion parameter had a Gaussian prior of *σ* = 50 ± 50. To reduce the stochasticity in the likelihood stemming from the non-deterministic nature of the IBM, all counts were spatially aggregated to larger cells of (25 km)^2^ and twenty IBM replicate runs were averaged for each sample. Posterior distributions were estimated using Markov chain Monte Carlo (MCMC) sampling. The sampler was a differential evolution sampler with memory and snooker updater (DEzs; Braak & Vrugt [45]) implemented by the R package BayesianTools [46]. The MCMCs converged for 8 out of the 9 selected species. Based on this, the goldcrest was excluded from further analysis.

### 2.3 Evaluation and analyses

To validate and evaluate model predictions, we examined both the full IBM simulations as well as the extracted DCRs separately. All evaluations were based on a sample of 200 draws taken from the joint posterior of each species.

The full simulation provides spatio-temporal projections of abundance, which were used to validate the model fit with RMSE (root mean squared error; Table A.3) and Harrell’s c-index (Newson [47]; Table 2) as well as by comparison with the Swiss breeding bird index [48]. The c-index is a rank correlation index that generalises the AUC index to non-binary response variables and we used its implementation in the Hmisc R-package. It quantifies the probability that the ranking of a given pair of model-projected abundances matches the respective ranking of MHB observations. To be able to compare to other distribution models, we also computed the AUC using occupancy probabilities derived from abundance predictions (Table A.3). For further validation, we generated model projections of total abundance time-series relative to the year 1999 (Fig. 1). They were compared to the Swiss breeding bird index, which directly estimates the same quantity from MHB data and was thus considered as reference (Fig. 1).

**Figure 1:**
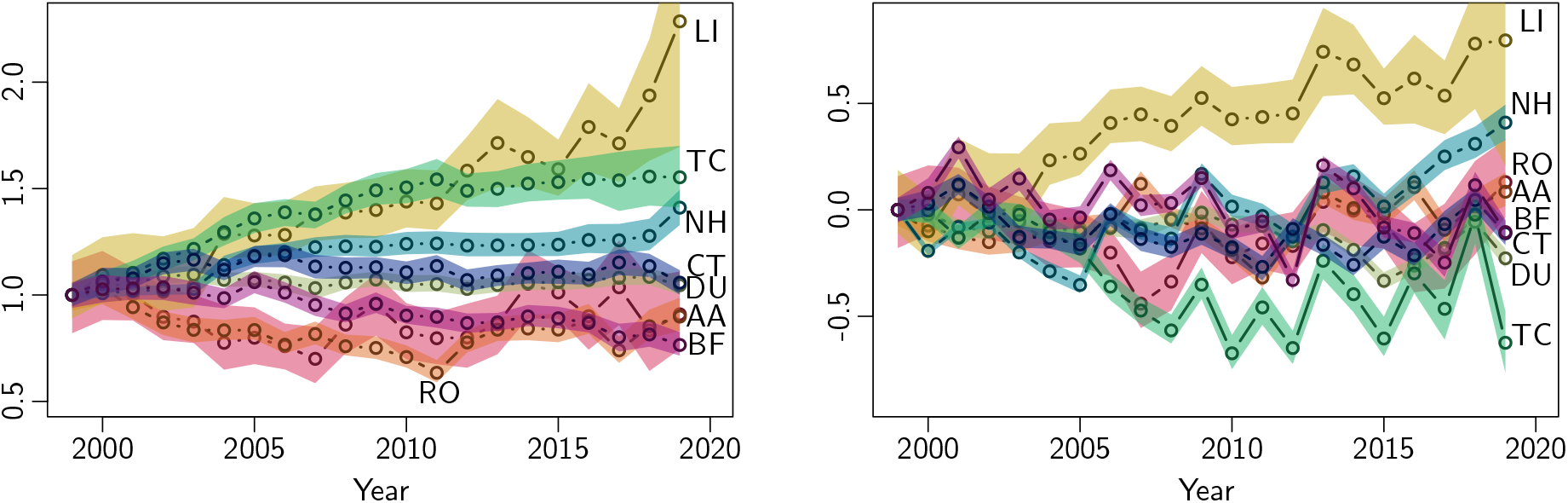
Total relative abundance time series (left) and its deviation from the Swiss breeding bird index (right), with 1999 (the first year of MHB data) as reference year. Shown are the median and the 95%-credible interval. BF - Bullfinch (pink); CT - Crested tit (dark blue); TC - Eurasian treecreeper (dark green); NH - Eurasian nuthatch (light blue); DU - Dunnock (light green); LI - Common linnet (yellow); RO - Ring ouzel (orange); AA - Alpine accentor (red).

**Table 2:** Model evaluation results for each species (column 1). The c-index (a rank correlation index) measures of goodness-of-fit (column 2). Column 3 states the low-density growth rate at median predictor values and columns 4 to 8 give its partial response to the observed trend in each climate variable. Details are given in the methods section. Colours code for strength of response, where grey denote little to no effect (less than 5% change), light red/blue denote small decrease/increase (more than 5% change), and dark red/blue denote strong decrease/increase (more than 10% change) in growth rate. The last column states the ratio of total simulated abundance under scenarios of observed and missing climate change. Given are the median and 95%-credible interval.

| | c-index | $r_{med}$ | $T_{br}$ | $P_{br}$ | $T_{at}$ | $T_{wn}$ | $P_{wn}$ | abund. change |
| --- | --- | --- | --- | --- | --- | --- | --- | --- |
| absolute change |  |  | +1.0 K | -11.6 mm | +1.1 K | +1.5 K | +29 mm |  |
| Bullfinch | 0.80 | 1.24 | 0.91 | 1.00 | 0.96 | 1.02 | 0.92 | 0.85 (0.83-0.90) |
| Crested tit | 0.75 | 0.99 | 0.92 | 0.97 | 0.97 | 1.01 | 0.97 | 0.80 (0.79-0.81) |
| E. treecreeper | 0.77 | 0.84 | 1.01 | 1.00 | 0.97 | 1.01 | 0.99 | 0.94 (0.92-0.99) |
| E. nuthatch | 0.74 | 0.94 | 1.09 | 1.05 | 1.00 | 1.03 | 1.12 | 1.08 (1.08-1.08) |
| Dunnock | 0.73 | 1.25 | 0.96 | 0.90 | 0.96 | 1.00 | 1.02 | 0.90 (0.89-0.90) |
| Common linnet | 0.73 | 1.36 | 1.01 | 1.05 | 0.98 | 1.01 | 1.11 | 1.18 (1.12-1.19) |
| Ring ouzel | 0.82 | 1.05 | 0.95 | 1.01 | 1.07 | 1.03 | 1.21 | 1.12 (1.10-1.22) |
| Alpine accentor | 0.88 | 1.04 | 0.94 | 0.96 | 0.98 | 1.01 | 1.09 | 1.12 (1.09-1.14) |

As a second step, we visually inspected the conditional response curves of all six DCRs within the range of observed climatic predictors for each species. This range spanned the 10%- and the 90%-quantiles of climate values occupied by a species, while the respective second predictor is held constant at its median. We then extracted the median and the 80% credible interval for each demographic rate, as shown in Fig. 2. The relationship for fecundity included density-dependence and thus had an additional parameter (1/*b*), describing the strength of density-dependence, and two additional predictors (local habitat suitability *h_i_* and density *n_i_*). For deriving the response curves, *h_i_* was held constant at its species-specific median and *n_i_* was set to one breeding pair per 1-km^2^ cell, yielding the fecundity that is realised at low densities.

**Figure 2:**
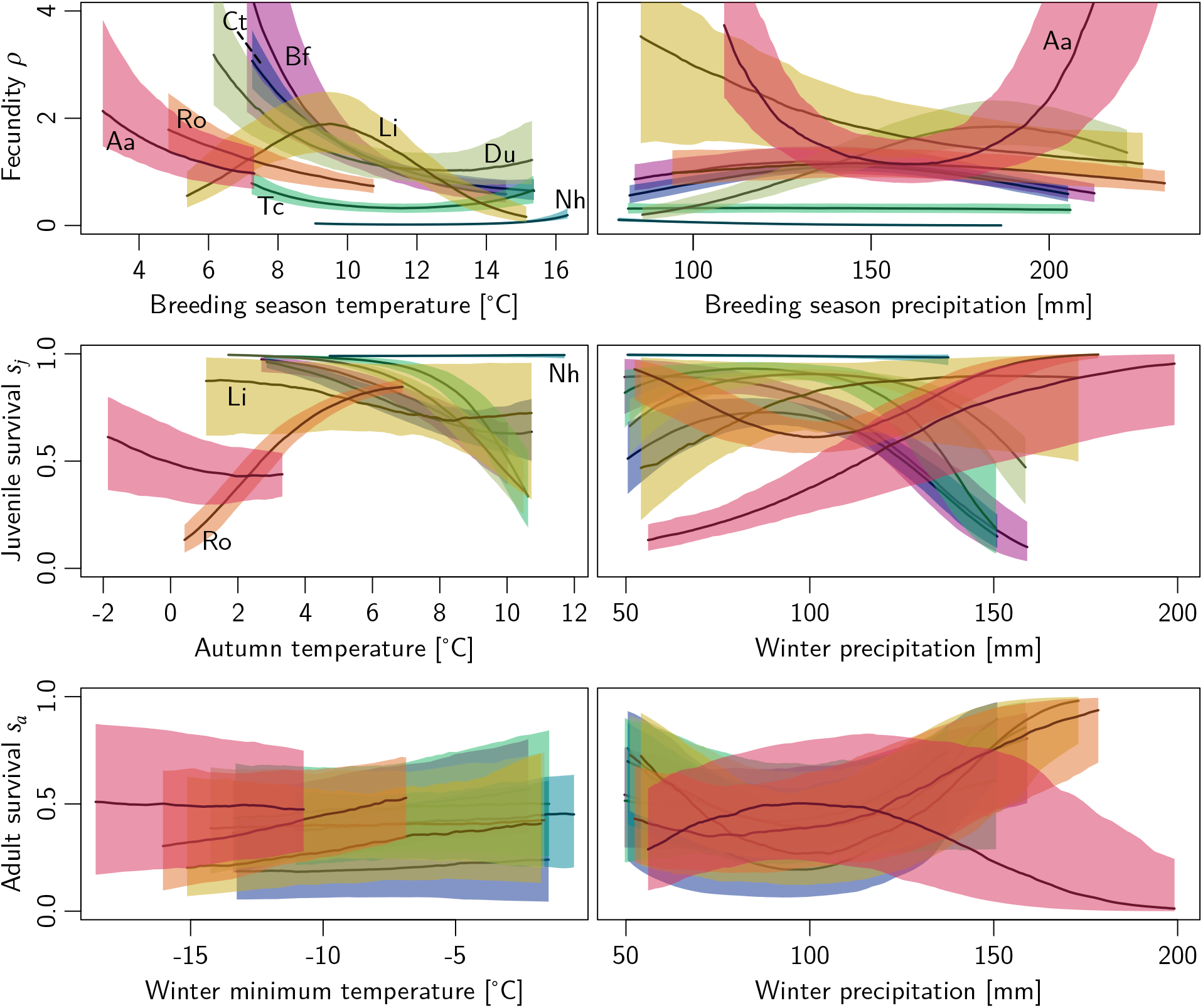
Demography-climate relationships for all species, evaluated over their respective typical climatic ranges. Shown are the median and 80%-credible interval. BF - Bullfinch (pink); CT - Crested tit (dark blue); TC - Eurasian treecreeper (dark green); NH - Eurasian nuthatch (light blue); DU - Dunnock (light green); LI - Common linnet (yellow); RO - Ring ouzel (orange); AA - Alpine accentor (red).

Lastly, we aimed to quantify the effects on predicted population performance that could be attributed to climatic trends over the observed time period. From the three demographic rates, we derived a local, low-density growth rate *r* as an overall measure of population performance. It was given by the leading eigenvalue of the transition matrix (as the population-based equivalent to our IBM), 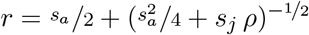. As a measure of vulnerability to climate change, we assessed the response of *r* to the observed changes in each climate variable individually (Table 2). For this, we compared a base growth rate, given as the value predicted at the median value of all predictors, with those for which a single predictor was changed. The amount of considered climatic change was determined from the linear trend in each predictor, accumulated over the twenty years of survey data (1999-2019). Thus, the vulnerability indicator combines the effects from both the sensitivity of the growth rate to climate change and the amount of exposure. However, it takes into account only the isolated effect from one predictor, while there are likely interactions between the impacts of different aspects of climate change. As an overall indicator of climate change impact, that simultaneously considers all predictors as well as their interactions, we used the full IBM to generate abundance projections under two scenarios: The factual, observed climate scenario and a counter-factual no-climate-change scenario, that consisted of trend-corrected predictors to simulate a stationary climate. The impact measure was then given by the ratio of projected mean total abundance over the years 2017-2019 under the scenarios of changing versus stationary climate.

## 3 Results

The goodness of fit of the posterior predictions was assessed using the c-index and RMSE for all species. The c-index ranged from 73% to 88% (Table 2), indicating acceptable to excellent model fit depending on the species. The RSME ranged from 1.2 to 5.7 and ranged models in a different order (Table A.3). In both measures, however, the model for alpine accentor scored best. The AUC consistently scored slightly above the c-index and ranged from 75% to 90% (Table A.3). This was expected because the c-index also reflects the accuracy of predicted abundances, not only occurrences. The IBM projections of total Swiss abundance are shown in Fig. 1, relative to the initial year when the MHB was launched (1999). For three species (linnet, treecreeper, nuthatch) the models predicted increasing trends, while stable or slightly declining trends were predicted for the remainder. However, these three models performed poorest in the validation and are therefore most uncertain. They scored lowest in terms of c-index and also showed the largest deviations from the Swiss breeding bird index, that indicated under-predicted abundance for the treecreeper and over-predicted abundances for linnet and nuthatch (Fig. 1).

The conditional response plots of all six demography-climate relationships (DCRs) are shown in Fig. 2, with the three response rates in rows and their respective temperature and precipitation predictors in columns. For each species, the DCRs were evaluated over their core occupied climatic range (given as the central 80% quantiles), while the respective second predictor is held constant at its median. The fecundity-temperature DCR indicated a trend of lower fecundity with increasing breeding season temperatures. Most species showed a monotonically decreasing relationship, though for the treecreeper and nuthatch it was almost constant, and only the linnet exhibited a clear optimum, at around 9°C. The fecundity-precipitation DCR was less certain, but showed a weak optimum for bullfinch, crested tit and ring ouzel, with maxima around 150 mm, and a pronounced optimum for the dunnock around 190 mm. Surprisingly, for the alpine accentor this relationship was bimodal (two optima at the boundaries of a steep inverted parabola), which is not ecologically plausible. The DCRs for juvenile survival consistently exhibited high values around 6°C mean autumn temperature for all species apart from the alpine accentor and and around 100 mm total winter precipitation for all species apart from alpine accentor and linnet. Individually, there were strong responses of juvenile survival to winter precipitation for the bullfinch (monotonous decrease), crested tit, treecreeper, dunnock, (unimodal) and alpine accentor (monotonous increase). The DCRs for adult survival were overall weaker and more uncertain than those for fecundity and juvenile survival. The relationship between adult survival and minimum winter temperature was fitted with only a first-order polynomial (Table 1) and results indicated a weak but consistent positive trend. Only the alpine accentor, a typical mountain bird, exhibited a constant relationship with increasing temperatures, which may be attributed to its adaptation to low temperatures. The DCR of adult survival with winter precipitation showed a minimum at around 100 mm for all species but the alpine accentor, that exhibited a maximum there. Some of the fitted DCRs have the shape of a parabola that opens upward, indicating optima at the extremes of their respective climatic range. We discuss possible explanations for these ecologically implausible DCRs and give remarks on their interpretation below.

To better understand the interplay of the different demographic processes, we summarised the three demographic rates obtained from the DCRs to a density-dependent growth rate *r.* Fig. 3 shows exemplary maps of all three demographic rates individually, and the resulting growth rate at low density (i.e., at one breeding pair per 1 km^2^) for the ring ouzel in 2018. Species can persist only in regions where the climatic conditions allow for all demographic rates to be high enough. The map of growth rates, Fig. 3 (d), highlights areas of negative population growth (*r* < 1) and those of positive growth (*r* > 1), thus providing an assessment of demographically suitable areas. For example, in the northern Alps all three demographic rates largely coincided in exhibiting high values, which resulted in positive growth of ring ouzel. In contrast, some areas in the southern Alps showed high fecundity but low survival probabilities, resulting in a negative population growth.

**Figure 3:**
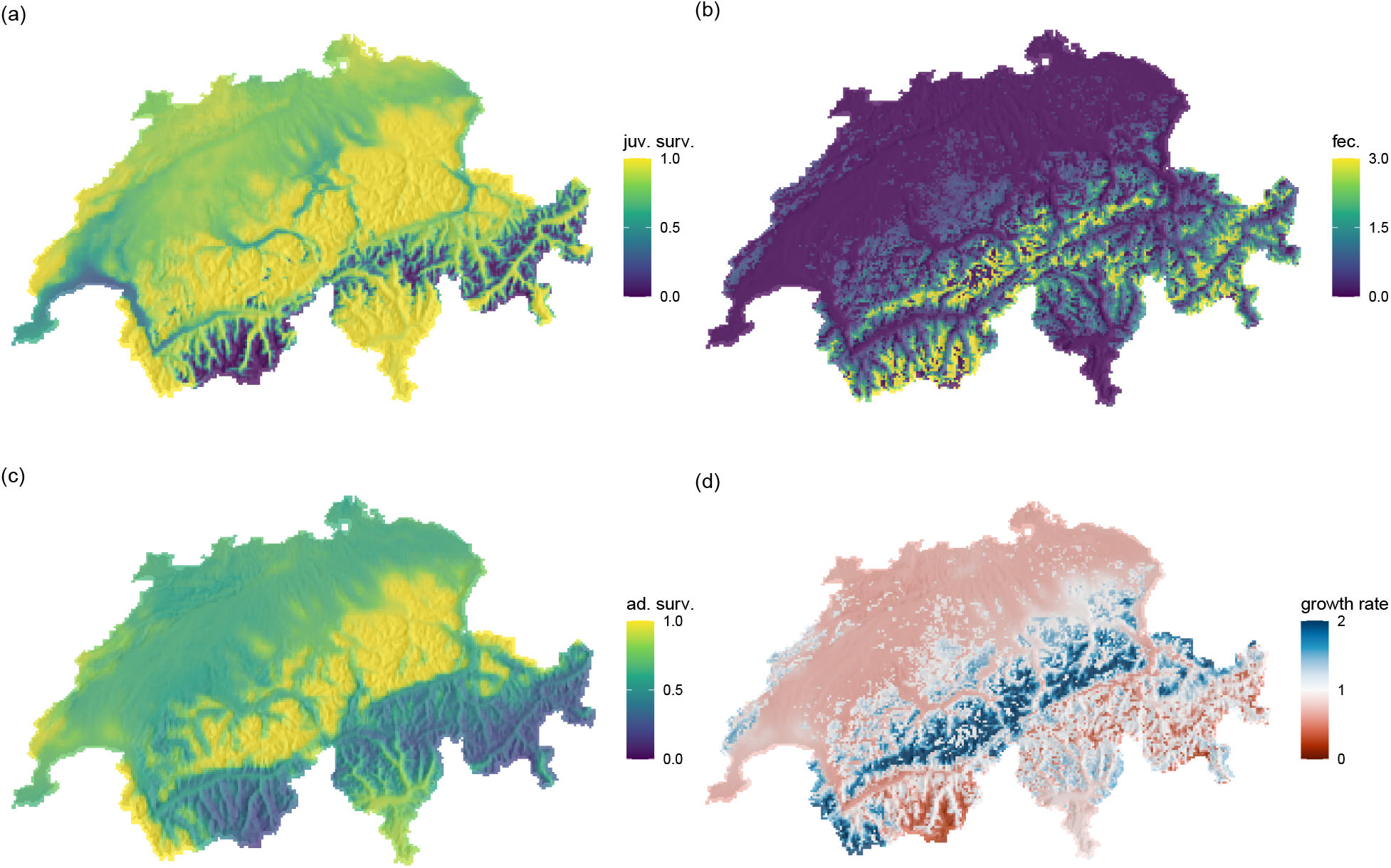
The three demographic rates juvenile survival (a), fecundity (b), adult survival (c), and the resulting growth rate at low densities (d) for the ring ouzel in year 2018. In (d), red/blue shades indicate growth rates below/above unity.

To assess the impact of the observed climate change on the study populations, we examined two measures of population response (Table 2): First, we calculated the change in low-density growth rate caused by the change of individual climate predictors that was observed over the survey period (second row in Table 2). According to this analysis, the most influential climate predictors were the mean temperature during breeding season, *T_br_*, which tended to decrease population growth, and precipitation during winter, *P_wn_,* which tended to increase population growth. To a lesser extent, breeding-season precipitation *P_br_* also caused significant but inconsistent changes. Some species were consistently impacted in one direction, e.g. the bullfinch and crested tit, that were adversely affected by both influential predictors, and the nuthatch, which benefitted from all three. Other species experienced mixed effects from the altered climate predictors on their growth rates, as alpine accentor and ring ouzel were adversely affected by the observed trends in *T_br_*, but gained from the changes in *P_wn_.* Secondly, we compared abundance predictions for the years 2017-2019 under the scenarios of observed climate versus no-climate change (last column in Table 2). According to this analysis, some species showed lower current abundances than would have been expected under a no-climate change scenario (bullfinch, crested tit, treecreeper, dunnock). The remaining species exhibited higher predicted current abundances (linnet, ring ouzel, alpine accentor) indicating that these species profited from climate change. Presumably, this benefit was primarily generated from increased winter precipitation. However, the predicted positive effect of climate change on abundance did not necessarily result in increasing populations. For example the ring ouzel and alpine accentor showed slight negative trends even under the more favourable scenario of observed climate change (Fig. 1).

## 4 Discussion

Spatio-temporal patterns of species abundance, as observed in monitoring or atlas surveys, are the result of a complex interplay of multiple ecological processes. In this study, we analysed the dynamics of Swiss breeding birds using individual- and process-based models that included local population dynamics and dispersal. Our models explicitly considered the demography-climate relationships (DCRs) of the constituent demographic processes juvenile survival, adult survival and fecundity. By simultaneously fitting all processes to survey data we were able to disentangle the effects of recent climate change on observed range and population dynamics. This allowed us to distinguish species that exhibited lower or higher abundance than predicted without climate change. Yet, results also indicated that care must be taken when interpreting the fitted DCRs as they need to be investigated closely for ecological plausibility. The calibration to time series of abundance data can impose trade-offs between the demographic responses, if model inadequacies like missing ecological processes or missing environmental predictors are present or the data contains insufficient information to separate the processes. In any case can the full model evaluations as well as the analysis of the individual DCRs improve our mechanistic understanding of range dynamics under climate change and pinpoint missing knowledge.

The increasing availability of long-term monitoring data allows fitting and validating complex process-based population models. In this study, we calibrated individual-based models (IBMs) that explictly include crucial processes such as density-dependence and climatic impacts on demographic rates [9]. To describe the climate, we used precipitation in addition to temperature as predictive variables [49]. When comparing our spatiotemporal predictions against observations, our models achieved acceptable to excellent predictive accuracy, with the highest performance for the alpine accentor. We also compared the total abundance time-series against the independently derived Swiss breeding bird index and found adequate fit of the temporal dynamics for most species. However, total abundance over time was markedly over-predicted for the linnet and nuthatch and under-predicted for the treecreeper. We can only hypothesise that this mismatch is due to violated model assumptions. Specifically, the heuristic habitat maps for the linnet may be overestimating the amount of suitable habitat. Based on published trait data for these species, the maps included a wide range of habitat classes (woodland, shrub, mountain meadows) while the true habitat preferences may be much more specific and met in only a smaller subset of the identified habitat areas. For nuthatch and treecreeper, their DCRs for fecundity were mostly constant in both predictors so that all spatial variation in fecundity stem from the habitat maps. For the nuthatch, also the juvenile survival was mostly constant. This suggests that important predictors for these two species may have been missing from the model. A recent study by Briscoe et al. [50] evaluated the predictive performance of dynamic occupancy models that represent local occurrence as a result from colonisation and extinction dynamics, but do not explicitly consider dispersal. Their models were fitted to the same MHB data for a large set of species and evaluated by AUC. In a direct comparison by AUC values our models mostly rank slightly lower, on average by five percentage points (Table A.3). However, our IBMs are expected to fit less closely to training data, as they use fewer spatial data (i.e. land use or climate predictors) and instead impose structure through specific assumptions on the generative, ecological processes.

The calibrated DCRs allow investigating the processes that underlie the spatial population dynamics in more detail. When interpreting the single demographic rates, it is important to consider their respective role in the model. Here, the survival rates include primarily winter and autumn mortality but do not account for dispersal mortality, which is represented by failed dispersal events and was not climate-dependent in our model. Thus, the effective annual survival is lower than given by the DCRs and the resulting local population dynamics alone. Further, fecundity described the number of independent juveniles produced per breeding pair and year. It therefore accounts for factors such as the number of broods per year and early juvenile mortality. Most of the DCRs show a monotonous or unimodal relationship, describing the relative favourability of climatic conditions and indicating optima. In the case of fecundity, above-median temperatures during breeding season tended to decrease it for all species. Juvenile survival attains very high values for some species under favourable climatic conditions, which can indicate high compensating dispersal mortalities. Adult survival tended to increase with winter temperature for all species. Thus, for some cases, general conclusions about climatic suitability across species can be drawn.

When interpreting the DCRs, however, care must be taken with respect to their transferability. All DCRs were calibrated to observed abundance data, which was modelled as the combined effect of the three demographic processes we consider. This can entail limited identifiability of parameters, if the same abundances are generated by different combinations of survival and fecundity. Further, if important ecological processes or environmental predictors are misrepresented or missing, the included processes will compensate for them, potentially leading to unexpected DCR shapes, that should be interpreted cautiously. For example, some of the fitted DCRs have the shape of inverted parabolas with two optima at the extremes of their range. This was found for the fecundity-precipitation DCR of the alpine accentor and the juvenile survival-precipitation DCR of the ring ouzel. Such a relationship is ecologically implausible as we would typically expect that demographic rates are optimal within certain climatic limits. A possible explanation for the alpine accentor is that the assumed fecundity-habitat relationship may not hold and the DCR attempts to compensate for low realised fecundities at the range margins, where low habitat suitability is projected. With respect to the ring ouzel, the local winter conditions may be poor predictors of survival since large parts of their Swiss population migrates during winter. For most species, the relationships of adult survival with winter precipitation was also slightly bimodal. This can be a result from interactions among adult survival and juvenile survival, which share minimum winter temperature as a common predictor. These two DCR tend to be negatively correlated for most species, apart from ring ouzel and alpine accentor, which supports this hypothesis. Such a negative correlation was also found by Dybala et al. [51], who modelled the impact of climate change on the demography of song sparrows in California. Tavecchia et al. [52] showed that climate-driven vital rates do not necessarily imply climate-driven population dynamics, especially in highly mobile species, as trade-offs can mask the changes in underlying processes. Further, an example for a potential missing mechanism are negative species interactions, whose presence can confound the value of an DCR, especially towards the range margins. Therefore, we advise that extrapolation to non-analogue climatic conditions outside the sampled range of training data should only be attempted if the fitted DCRs appear plausible. Despite these limitations in interpretability of the individual DCRs, their combined effects within the full model simulations were still able to reproduce the observed abundances and yield good model fit. Importantly, DCRs allow an improved mechanistic understanding of the underlying processes and potentially missing information, which is a substantial advantage over simple hybrid models [27, 28].

By combining the demographic rates to an overall growth rate, we can better understand the causes underlying the demographic suitability of different geographic regions. Further, the derivation of spatially and temporally explicit growth rates allows more standardised interpretations across different model frameworks. For example, we found that the ring ouzel was predicted to generally persist in high altitudes where fecundity is high enough, but that some of these high-altitude areas are climatically unsuitable due to high winter mortality. According to Barras et al. [39], future losses in abundance are expected especially in the Northern Alps, where we predicted highest current growth rates, and gains in abundance in the Central Alps, where we predicted lowest current growth rates. To assess the impacts of observed climatic changes on the demographic performance of the studied species, we considered two measures of climate vulnerability. First, evaluating the conditional impact of each climatic predictor identified breeding-season temperature and winter precipitation as the most influential variables. This impact is a combined effect of the magnitude of already observed climate change and the sensitivity of the growth rate to them. However, more targeted investigations are needed to understand the biological pathways by which a given abiotic factor controls an ecological process. For example, Barras et al. [53] explained for ring ouzel that elevated temperatures during breeding negatively impacted the feeding of chicks by parents. In fact, our model confirms a negative effect of breeding-season temperature on growth rate for this species. Also our modelled current growth rate (1.05) coincides with the one measured by Barras et al. [54] (1.04). Second, comparing the simulated total abundances under observed climate against a no-climate change scenario, we were able to estimate in how far the predicted abundance trends could be attributed to climate change. Such analyses are useful to understand the overall direction of the combined effects of climate change on populations. For example, ring ouzel and alpine accentor show slightly declining population trends, both in the breeding bird index and in our projections. However, our models indicated that predicted abundance was still higher then in the no-climate change scenario, implying that both species would experience even stronger population declines without recent climate change. Overall, our models suggested that typical mountain species tend to benefit from current climate change (without necessarily amounting to positive population trends), while lowland species are already negatively affected by climate change. We do not expect the trend for mountain species to extend far into the future, when climatic changes become more pronounced. In fact, while we assessed the effect of contemporary climate change only, the vulnerability of Swiss breeding birds to future climate change was investigated previously [55], by comparing current climatic conditions with those in 2100. Within this time frame, alpine accentor and ring ouzel are expected to suffer strong range contractions. Here, correctly identified DCRs can help to identify promising avenues for effective management. By explicitly investigating how individual demographic rates are affected by climate change, targeted measurements can be designed to support these vulnerable processes and mitigate climate change effects.

Considering the strong responses of species distributions to recent climatic changes, it becomes clear that a mechanistic understanding of the underlying processes is needed. The calibration of detailed, process-based models from data that emerged under a range of climatic conditions allows to disentangle the different processes that drive spatial population dynamics and understanding the differential effects of climate change. The estimation of DCRs in this study provided a better mechanistic understanding of the responses of eight (of nine) Swiss breeding birds to contemporary climate change. In a next step, more abiotic and biotic mechanisms should be tested, for example by extending the set of regarded environmental predictors and by considering different alternative model structures. It is important to acknowledge that the model set-up still relies on inverse calibration and is thus phenomenological to a certain extent. The resulting DCRs need to be interpreted cautiously and inspected for plausibility before drawing conclusions for targeted conservation management. This also requires profound insights into the biological mechanisms, by which ecological processes are influenced by the abiotic and biotic environment. For future research, we advocate a stronger integration of direct measurements of DCRs, where this is feasible, with inverse calibration and verification from independent abundance time series. Ultimately, this will increase our confidence for drawing conclusions on the mechanisms underlying complex range and population dynamics and making predictions into the future.

## Acknowledgements

We thank Guillermo Fandos, Arnaud Barras, Pius Korner, Michael Schaub, and Nicolas Strebel for valuable comments. AM and DZ were supported by Deutsche Forschungsgemeinschaft (DFG) under grant agreement No. ZU 361/1-1.

## Author Contributions

A.-K. Malchow: Conceptualization (equal), Formal Analysis (lead), Methodology (equal), Software (lead), Data Curation (support), Visualization (lead), Writing – Original Draft Preparation (lead), Writing – Review & Editing (equal). F. Hartig: Conceptualization (support), Formal Analysis (support), Methodology (equal), Supervision (support), Writing – Review & Editing (equal). J. Reeg: Software (support), Visualization (support), Writing – Review & Editing (equal). M. Kéry: Data Curation (lead), Writing – Review & Editing (equal). D. Zurell: Conceptualization (equal), Methodology (equal), Funding acquisition (lead), Supervision (lead), Writing – Review & Editing (equal).

## Data Availability

The study uses data from the Swiss breeding bird survey and the Swiss breeding bird index provided by the Swiss ornithological institute, Sempach. All scripts and data required to run the presented analyses can be accessed from the public GitHub repository: https://github.com/UP-macroecology/Malchow_DemogEnv_2022. Upon acceptance, we will create a permanent Zenodo archive and provide its DOI. The utilised R-packages are open-source software: RangeShiftR is available from GitHub under https://github.com/RangeShifter/RangeShiftR-package. BayesianTools is available directly from CRAN or from GitHub under https://github.com/florianhartig/BayesianTools.

## A Supplemental information

**Table A.1:** Mean (〈·〉) and standard deviation (SD; *σ*) of the Gaussian prior distributions for demographic and dispersal parameters for each species, derived from [42]. Columns: fecundity *ρ* and its SD *σ_ρ_*; juvenile survival probability *s_j_*, adult survival probability *s_a_,* emigration probability *p_e_,* and their SDs *σ_s,p_*; mean dispersal distance 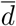 and its SD 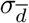 given in kilometres. The prior of overall demographic density dependence is given by 1/*b* = 0.003 ± 0.002 for all species. *ρ*, *s_j_*, and *s_a_* denote the intercepts of their respective demography-climate relationships.

| Species | $\langle \rho \rangle$ | $\sigma_\rho$ | $\langle s_j \rangle$ | $\langle s_a \rangle$ | $\langle p_e \rangle$ | $\sigma_{s,p}$ | $\langle \bar{d} \rangle$ | $\sigma_{\bar{d}}$ |
| --- | --- | --- | --- | --- | --- | --- | --- | --- |
| Bullfinch | 2.25 | 0.50 | 0.50 | 0.47 | 0.37 | 0.20 | 1.36 | 0.50 |
| Crested tit | 1.62 | 0.06 | 0.50 | 0.27 | 0.50 | 0.20 | 1.44 | 0.50 |
| Eurasian treecreeper | 2.06 | 0.56 | 0.50 | 0.50 | 0.42 | 0.20 | 1.37 | 0.50 |
| Eurasian nuthatch | 1.75 | 0.44 | 0.50 | 0.45 | 0.16 | 0.20 | 1.69 | 0.50 |
| Goldcrest | 5.00 | 0.88 | 0.50 | 0.50 | 0.52 | 0.20 | 1.48 | 0.50 |
| Dunnock | 2.50 | 0.50 | 0.50 | 0.49 | 0.79 | 0.20 | 2.66 | 0.50 |
| Common linnet | 3.12 | 0.62 | 0.50 | 0.38 | 0.51 | 0.20 | 1.46 | 0.50 |
| Ring ouzel | 1.50 | 0.28 | 0.38 | 0.52 | 0.09 | 0.20 | 2.38 | 0.50 |
| Alpine accentor | 1.75 | 0.38 | 0.50 | 0.50 | 0.45 | 0.20 | 1.39 | 0.50 |

**Table A.2:** Species habitat preferences reflected in habitat maps. Habitat types from [42]: Deciduous forest (Dec. F.), Coniferous forest (Con. F.), Woodland (Wdl.), Shrub, Grassland (Grl.), Mountain meadows (M. mead.), Rocks. Indices of CORINE classes that were mapped from habitat types: 231 - Pastures, 311 - Broad-leaved forest, 312 - Coniferous forest, 313 - Mixed forest, 321 - Natural grasslands, 322 - Moors and heathland, 324 - Transitional woodland-shrub, 332 - Bare rocks, 333 - Sparsely vegetated areas.

| Name english | Habitat preference | Corine classes |
| --- | --- | --- |
| Bullfinch | Dec. F., Con. F. | 311, 312, 313 |
| Crested tit | Dec. F., Con. F. | 311, 312, 313 |
| Eurasian treecreeper | Dec. F., Con. F. | 311, 312, 313 |
| Eurasian nuthatch | Dec. F., Con. F. | 311, 312, 313 |
| Dunnock | Dec. F., Con. F., Wdl., Shrub | 311, 312, 313, 321, 324 |
| Common linnet | Wdl., Shrub, M. mead. | 231, 321, 322, 324, 333 |
| Ring ouzel | Con. F., Grl., Rocks | 312, 322, 332, 333 |
| Alpine accentor | M. mead., Rocks | 322, 332, 333 |

**Table A.3:** Model performance as measured by RSME and comparison of our IBMs and the dynamic occupancy models presented in Briscoe et al. [50]. The comparison is done by AUC, where the abundances predicted by our IBMs were compressed to presence-absence data (P/A).

| Species | RSME | AUC of P/A | AUC from Briscoe et al. (2021) |
| --- | --- | --- | --- |
| Bullfinch | 2.4 | 0.84 | 0.88 |
| Crested tit | 4.8 | 0.79 | 0.86 |
| E. treecreeper | 3.4 | 0.83 | 0.87 |
| E. nuthatch | 4.2 | 0.82 | 0.89 |
| Dunnock | 5.7 | 0.77 | 0.85 |
| Common linnet | 1.9 | 0.75 | 0.77 |
| Ring ouzel | 3.5 | 0.86 | 0.94 |
| Alpine accentor | 1.2 | 0.90 | — |

